# Dual function for Tango1 in secretion of bulky cargo and in ER-Golgi morphology

**DOI:** 10.1101/144923

**Authors:** LD Rios-Barrera, S Sigurbjörnsdóttir, M Baer, M Leptin

**Affiliations:** European Molecular Biology Laboratory, 69117 Heidelberg, Germany; Institute of Genetics, University of Cologne, 50674 Cologne, Germany; Current address: University of Iceland, 101 Reykjavík, Iceland; Current address: Ludwig-Maximilian University of Munich, 81377 Munich, Germany

**Keywords:** Tango1, ER/Golgi, secretion, Drosophila

## Abstract

Tango1 helps the efficient delivery of large proteins to the cell surface. We show here that loss of Tango1, in addition to interfering with protein secretion, causes ER stress and defects in cell and ER/Golgi morphology. We find that the previously observed dependence of smaller cargos on Tango1 is a secondary effect, due to an indirect requirement: if large cargos like Dumpy, which we identify here as a new Tango1 cargo, are removed from the cell, non-bulky proteins re-enter the secretory pathway. Removal of the blocking cargo also attenuates the ER-stress response, and cell morphology is restored. Thus, failures in the secretion of non-bulky proteins, ER stress and defective cell morphology are secondary consequences of the retention of cargo. By contrast, the ERES defects in Tango1-depleted cells persist in the absence of bulky cargo, showing that they are due to a secretion-independent function of Tango1. Therefore, the maintenance of proper ERES architecture may be a primary function for Tango1.

## Introduction

The endoplasmic reticulum (ER) serves as a major factory for protein and lipid synthesis. Proteins and lipoproteins produced in the ER are packed into COPII-coated vesicles, which bud off at ER exit sites (ERES) and then move towards the Golgi complex where they are sorted to their final destinations. Regular COPII vesicles are 60 - 90 nm in size, which is sufficient to contain most membrane and secreted molecules (Szul and Sztul, 2011). The loading of larger cargo requires specialized machinery that allows the formation of bigger vesicles to accommodate these bulky molecules. Tango1 (Transport and Golgi organization 1), a member of the MIA/cTAGE (melanoma inhibitory activity/cutaneous T cell lymphoma-associated antigen) family, is a key component in the loading of such large molecules into COPII-coated vesicles. Molecules like collagens and ApoB (apolipoprotein B)-containing chylomicrons are 250-450 nm long and rely on Tango1 for their transport out of the ER, by physically interacting with Tango1 or Tango1 mediators at the ERES (Pfeffer, 2016; Saito et al., 2009; Santos et al., 2016).

Tango1 is an ER transmembrane protein that orchestrates the loading of its cargo into vesicles by interacting with it in the ER lumen. The interaction of Tango1 with its cargo then promotes the recruitment of Sec23 and Sec24 coatomers on the cytoplasmic side, while it slows the binding of the outer layer coat proteins Sec13 and Sec31 to the budding vesicle. This delays the budding of the COPII carrier (Saito et al., 2009). Tango1 also recruits additional membrane material to the ERES from the Golgi intermediate compartment (ERGIC) pool, thereby allowing vesicles to grow larger (Santos et al., 2015). It also interacts directly with Sec16, which is proposed to enhance cargo secretion (Maeda et al., 2017).

Apart from bulky proteins, some heterologous, smaller proteins like secreted horseradish peroxidase (ssHRP, 44 kDa) and secreted GFP (27 kDa) also depend on Tango1 for their secretion (Nogueira et al., 2014). Unlike for collagen or ApoB, there is no evidence for a direct interaction between Tango1 and ssHRP or secreted GFP. It is not clear why Tango1 would regulate the secretion of these molecules, but it has been proposed that in the absence of Tango1, the accumulation of non-bulky proteins at the ER might be due to abnormally accumulated Tango1 cargo clogging the ER (Nogueira et al., 2014; Saito et al., 2009); however, this has not been tested experimentally.

Drosophila Tango1 is the only member of the MIA/cTAGE family found in the fruit fly, which simplifies functional studies. Like vertebrate Tango1, the Drosophila protein participates in the secretion of collagen (Lerner et al., 2013; Pastor-Pareja and Xu, 2011). And as in vertebrates, ssHRP, secreted GFP and other non-bulky molecules like Hedgehog-GFP also accumulate in the absence of Tango1 (Bard et al., 2006; Liu et al., 2017). These results have led to the proposal that Tango1 participates in general secretion. However, most of the evidence for these conclusions comes from overexpression and heterologous systems that might not reflect the physiological situation.

Here, we describe a *tango1* mutant allele that we identified in a mutagenesis screen for genes affecting the structure and shape of terminal cells of the Drosophila tracheal system (Baer et al., 2007). Tracheal terminal cells form highly ramified structures with branches of more than 100 mm in length that transport oxygen through subcellular tubes formed by the apical plasma membrane. Their growth relies heavily on membrane and protein trafficking, making them a very suitable model to study subcellular transport. We used terminal cells to study the function of Tango1, and we found that loss of Tango1 affects general protein secretion indirectly, and it also leads to defects in cell morphology and in the structure of the ER and Golgi. The defects in ER and Golgi organization of cells lacking Tango1 persist even in the absence of Tango1 cargo.

We identify a new bulky cargo for Tango1 in Drosophila. Our studies have allowed us to explain why in the absence of Tango1, non-bulky proteins accumulate in the ER in spite of not being direct Tango1 cargos. We show that these cargos are retained in the ER as a consequence of non-secreted bulky proteins interfering with their transport. However, the effect of loss of Tango1 on ER/Golgi morphology can be uncoupled from its role in bulky cargo secretion.

## Materials and Methods

### Fly stocks and genetics

All experiments were done at 25°C in standard conditions. To generate homozygous mutant *tango1* terminal cells we used the MARCM system (Baer et al., 2007), with the lines *hsFlp1.22; tub-GAL80, FRT40A; btl-GAL4, UAS-eGFP* (from Stefan Luschnig, University of Muenster, Germany), and the line *FRT40A* as control (Bloomington Drosophila Stock Center [BDSC] #5615). *2L3443* was mapped by complementation tests with *Df(2L)BSC7* (BDSC #6374), *Df(2L)BSC6* (BDSC #6338) and *Df(2L)BSC187* (BDSC #9672), followed by fine mapping through ORF sequencing of the genes within the segment genetically defined to contain the mutation. Final complementation tests with *tango1^GS17108^* (Kyoto Drosophila Genetic Resource Center [DGRC] #206906), and *tango1^GS15095^* (DGRC #206078) confirmed 2L3443 as a *tango1* allele.

The lines used as drivers for UAS constructs were *SRF-gal4* (Guillemin et al., 2001), *Lpp-gal4* (Palm et al., 2012), *nub-gal4* (Pastor-Pareja, and Xiu, 2014, #63148), *repo-gal4* (from Christian Klämbt, University of Münster, Germany), and *sr-gal4* (from Frank Schnorrer, Developmental Biology Institute of Marseille (IBDM), France). The following lines were obtained from Vienna Drosophila Resource Center: *ergic53^fTRG^* (#318063), *lanA^fTRG^* (#318155), *lanB1^fTRG^* (#318180) and *BM-40-SPARC^fTRG^* (#318015), which are fosmid constructs expressing GFP fusion proteins at endogenous levels (Sarov et al., 2016), and *UAS-pio-IR* (#107534). The *UAS-tango1-IR* (#11098R-3) and *UAS-vkg-IR* (#16858R-1) were obtained from the National Institute of Genetics Fly Stock Center, Japan. *dpy-YFP* and *UAS-dpy-IR* are from Barry Thompson, The Francis Crick Institute, UK (Ray et al., 2015). Collagen-GFP is a protein trap insertion of GFP in the *vkg* locus resulting in a fusion of collagen and GFP (Morin et al., 2001). *UAS-crb^extraTM^-GFP* is a construct where the cytoplasmic end of Crb was replaced by GFP (Pellikka et al., 2002). *UAS-Gasp-GFP* is from Christos Samakovlis, Stockholm University, Sweden (Tiklova et al., 2013). *UAS-Xbp1-GFP* is from Pedro Domingos, Nova University of Lisbon, Portugal (Ryoo et al., 2007). The following lines were obtained from BDSC: *UAS-ManII-GFP* (#65248), *UAS-RFP-KDEL* (#30910 and #30909), *UAS-mCD8mCherry* (#27392), *UAS-myrRFP* (#63148). *UAS-βPS-Integrin-Venus* was generated by subcloning *βPS-Integrin-Venus* from *pUbi-βPS-Integrin-Venus* [from Guy Tanentzapf, University of British Columbia, Canada (Yuan et al., 2010)] into the *pUASTattB* vector and then inserting in the third chromosome (VK33, BDSC #9750). *UAS-tango1-GFP* was generated by cloning the full-length *tango1* cDNA (GH02877) into pDONR221 (Gateway System, Invitrogen). This was recombined into the destination vectors pTWG from the Drosophila Gateway Vector Collection using the Gateway LR reaction. The construct was then subcloned into *pUASTattB* and injected into VK33.

### Whole mount sample preparation, microscopy and analyses

For tracheal terminal cell analyses, third instar wandering larvae were heat-fixed in Halocarbon oil for 30 seconds at 65°C. For tendon cell analyses, pupae at 24h after puparium formation were hand-peeled and immobilized with heptane glue in MatTek plates with Halocarbon oil. In both cases, samples were imaged immediately using a Zeiss LSM 780 confocal microscope. Quantitative analyses of the number of branching points and air-filling were performed in dorsal terminal cells in metameres 3-6 of heat-fixed larvae. Branches were counted manually in FIJI (Schindelin et al., 2012). Analysis of air-filling was performed by visualizing the presence of lumen using light transmission.

### Immunofluorescence staining

We used the following antibodies: guinea pig anti-Tango1 (1:400, from Sally Horne-Badovinac, University of Chicago, USA), rabbit anti-Sec16 (1:600, from Catherine Rabouille, Hubrecht Institute, Netherlands), rat anti-Crb (1:500, from Elisabeth Knust, MPI-CBG, Germany), rabbit anti-Pio [1:300, from Markus Affolter, University of Basel, Switzerland (Jazwinska et al., 2003)], mouse anti-βPS Integrin (1:200, DSHB #6G11), rabbit anti-Sec23 (1:200, Thermo Scientific #PA1-069), rabbit anti-GM130 (1:500, Abcam #ab30637), and rabbit anti-Dof [1:200 (Vincent et al., 1998)]. Alexa-conjugated antibodies from Thermo Scientific: Alexa568 goat anti-mouse (A-11031), Alexa568 goat anti-rat (A-11077), Alexa647 goat anti-rat (A-21247), Alexa568 goat anti-rabbit (A-11036), Alexa647 goat anti-rabbit (A-21245), Alexa568 goat anti-guinea pig (A-11075), Alexa647 goat anti-guinea pig (A-21450). Chromotek’s GFP-booster coupled to Atto488 (gba488) and RFP-booster coupled to Atto594 (rba594) were used to enhance signal from fluorescent reporters.

Third instar wandering larvae were collected, dissected, fixed using 4% PFA in PBS for 20 min and washed with PBTx (0.3% Triton X-100 in PBS) followed by 1 h incubation in blocking solution (PBTx, 1% BSA). Primary antibodies were diluted in blocking solution and incubated overnight at 4°C. After washing with PBTx, samples were incubated with secondary antibodies diluted in blocking solution at room temperature for 90 min followed by extensive washing using PBTx. Samples were mounted for imaging using Vectashield with DAPI (Vector Laboratories) and images acquired on Leica SP2, Zeiss LSM 780 or Zeiss LSM 880 Airyscan confocal microscopes.

### Western blotting

We used guinea pig anti-Tango1 (1:10,000, mentioned above) and mouse anti-βTubulin (1:5000, Amersham Life Science). HRP-conjugated antibodies were from Jackson ImmunoResearch Laboratories: goat anti-guinea pig-HRP (106-035-003) and goat anti-mouse-HRP (115-035-003). For each genotype, 20 embryos were selected, homogenized in loading buffer and heated for 5 min at 95°C. Samples were then separated by SDS-PAGE, transferred to PVDF membranes and subjected to immunodetection using the Luminata Crescendo Western HRP system.

### Image analyses

All analyses were done using FIJI. We determined the amount of collagen surrounding terminal cells by quantifying the fluorescence intensity of collagen-GFP at the cell membrane close to the terminal cell body. We subtracted the background from the mean fluorescence intensity of an area of 3 × 30 pixels within a single confocal plane. To determine Sec16 particle size and number, and Dpy and laminin accumulation, we masked the channel of interest with the contour of the cell or tissue of interest, and then segmented individual dots from maximum projection images. For Dpy accumulation in wing discs, we used the plot profile function. All images within an experiment were acquired using the same microscope settings.

### Statistical analyses

We used GraphPad Prism 6 for all statistical analyses. Plots were generated using GraphPad Prism 6 or Microsoft Excel.

## Results

### Identification of a mutation in *tango1*

Terminal cells of the tracheal system extend long subcellular branches that transport gas through tubes formed by the apical plasma membrane. The tubes can be easily visualized by bright field microscopy because of the difference in refractive index between the cytoplasm and the gas, providing a simple readout for branch maturation (Tsarouhas et al., 2007). In a screen for genes necessary for tracheal terminal cell branching, we identified a mutation, 2L3443, which caused air-filling defects and reduced branch numbers in homozygous mutant terminal cells [Fig. 1A-D (Baer et al., 2007)]. The mutation is embryonic semi-lethal (33.3% of homozygous embryos failed to hatch), and survivors died at early larval stages. We mapped this mutation by SNP recombination (Berger et al., 2001) and by complementation tests with deficiencies to the region 26D10-26F3 on the cytogenetic map (Fig. S1A). We identified a mutation within the ORF of *tango1*, and confirmed it is allelic to other *tango1* mutant alleles (Figure S1A’). The mutation introduces a premature stop codon in amino acid 1341 (Arginine to stop codon) downstream of the Proline-rich domain (PRD) that results in a truncation of the last 89 amino acids of the predicted protein (Fig. S1B, C). The missing segment contains an arginine-rich domain that has no predicted interaction partners. A Tango1-GFP construct expressed under a trachea-specific promoter suppressed the mutant phenotype (Fig. 1C-D) and an interfering RNA (tango1-IR) expressed specifically in terminal cells caused the same air-filling defects and reduction in branch number (Fig. 2A-B, E), confirming that *tango1* disruption was responsible for the branching defects.

**Figure 1.**
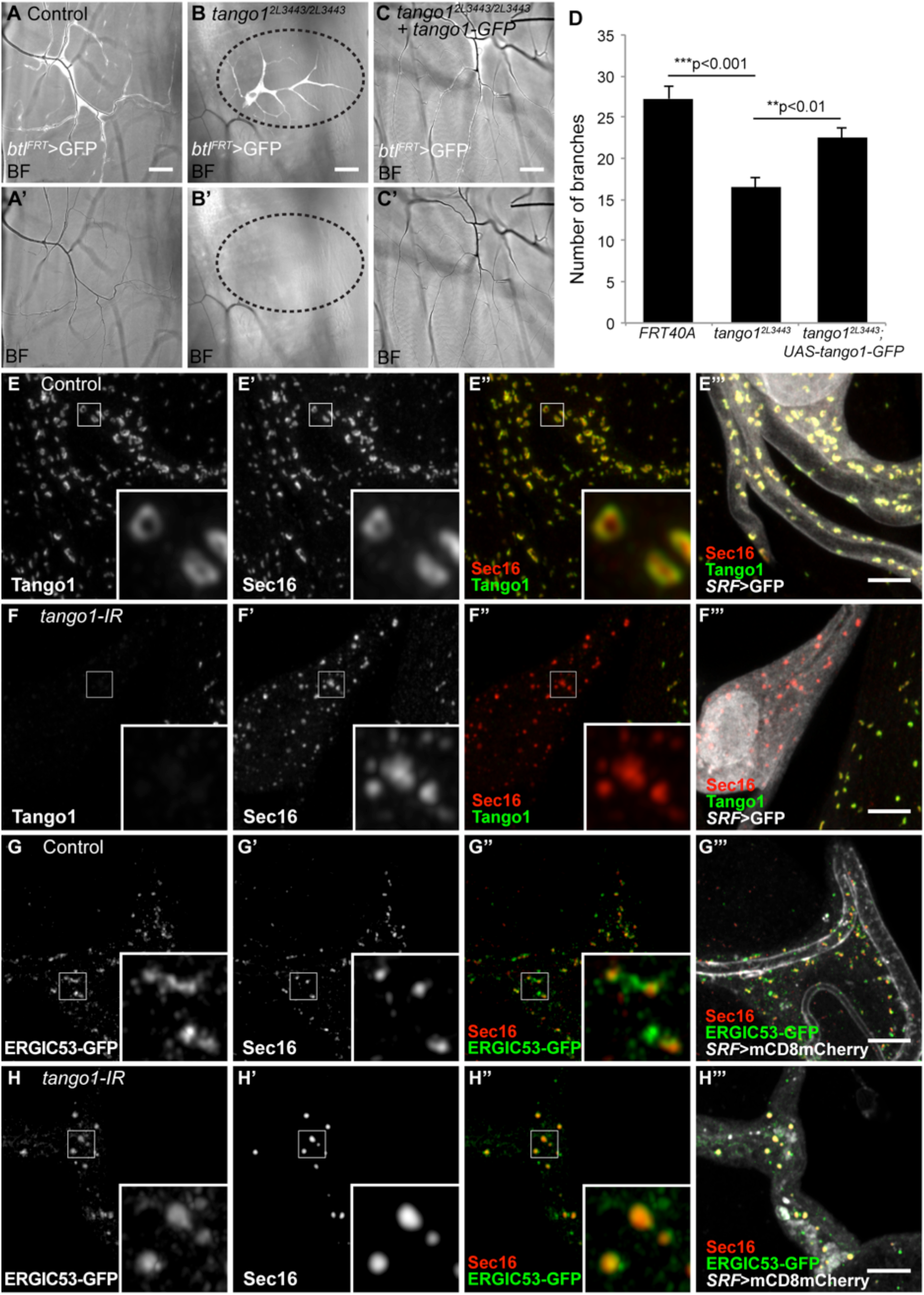
Effect of loss of Tango1 on cell, ER and Golgi morphology. (A-C) Bright field (BF) images of homozygous *tango1^2L3443^* mutant tracheal cells expressing GFP (btl^FRT^>GFP) allow the visualization of number of branches and the presence of air in terminal cells. Unlike control cells (A), homozygous *tango1^2L3443^* cells are not air-filled (area surrounded by dotted line in B). (C) Expression of Tango1-GFP in mutant cells suppresses the air-filling defects and re-establishes near-normal number of branches (D). Control, n=11; *tango1^2L3443^*, n=14; tango1^2L3443^+Tango1-GFP, n=11. Bars represent mean +/-SEM. Significance was determined using two-tailed t-test. (E-H) Airyscan microscopy images of control (E, G) and *tango1* knockdown cells (F, H), stained for Sec16 and Tango1 (E, F) and for ERGIC53-GFP (fTRG library, expressed at endogenous levels) and Tango1. Scale bars are 40μm (A-C) and 5μm (EH).

**Figure 2.**
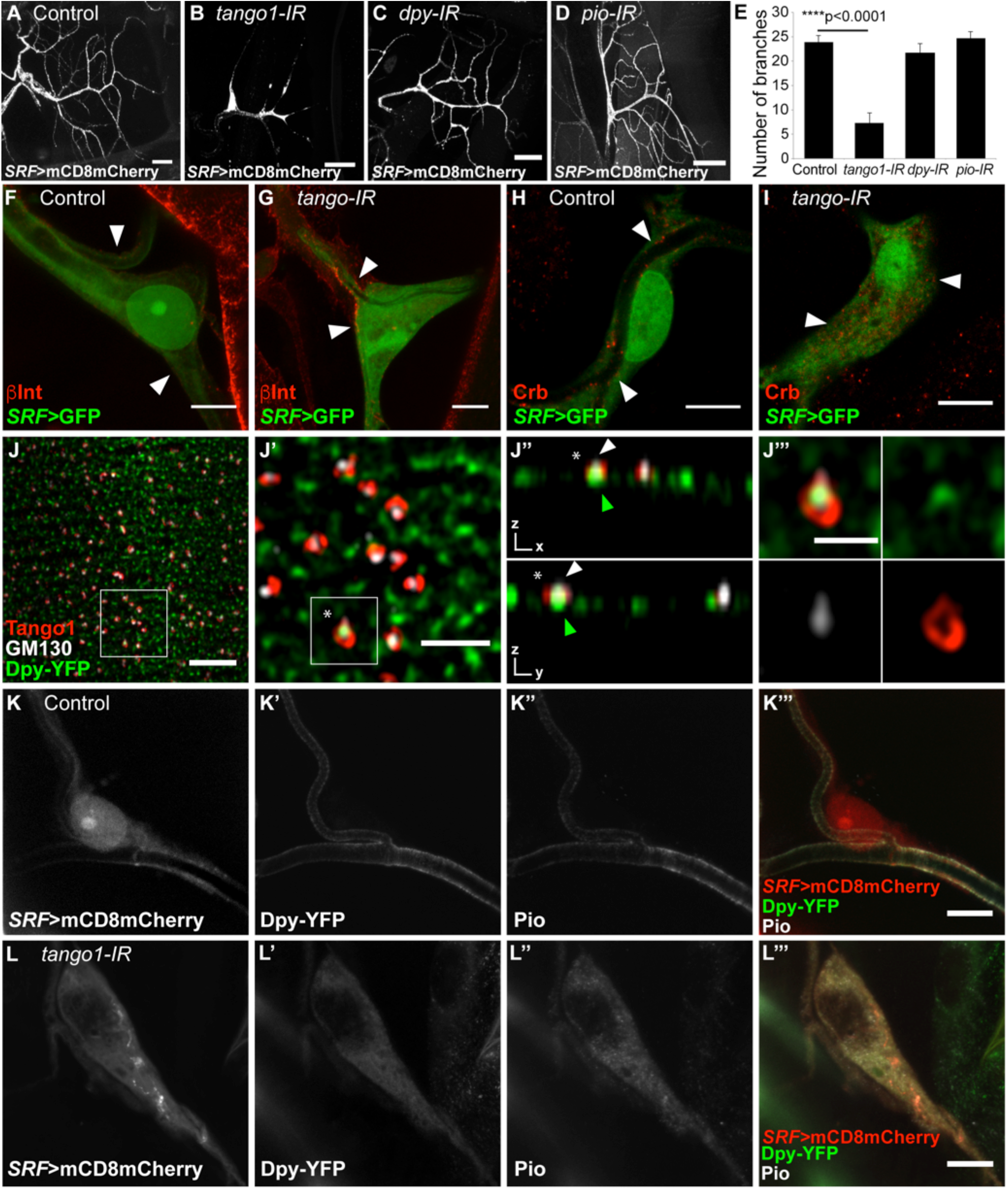
Function of cargo proteins and their distribution upon loss of Tango1. (A-E) Terminal cells were visualized by expressing mCD8mCherry under the terminal-specific driver *SRF-gal4*. (E) Manual quantification of branch numbers in terminal cells expressing different RNAi; cells expressing *tango1* RNAi (B) have fewer branches than control cells (A). Neither *dpy* RNAi (C) nor *pio* RNAi (D) affect branch numbers. Control, n=8; *tango1-IR*, n=9; *dpy-IR*, n=8; *pio-IR*, n=9. Bars represent mean +/− SEM. Significance was determined using two-tailed t-test. (F, G) Confocal projections of control (F) and *tango1-IR* (G) terminal cells expressing *SRF>GFP* and stained for βPS integrin (βInt). Arrowheads point to βInt localization. (H, I) Confocal projections of control (H) and *tango1-IR* (I) terminal cells expressing *SRF>GFP* and stained for Crb. Arrowheads point to Crb localization. (J) Airyscan images of details of tracheal dorsal trunk cells expressing Dpy-YFP and stained for Tango1 and Golgi marker GM130. White squares indicate the magnified regions. (J’’) orthogonal views of a single plane from (J’). (K, L) Confocal projections of control (K) and *tango1-IR* (L) terminal cells expressing SRF>mCD8mCherry and Dpy-YFP and stained for Pio. Scale bars are 40μm (A-D), 10μm (F-I, K-L), 5μm (J), 2μm (J’) and 1μm (J’’’).

To determine the role of Tango1 in terminal cells, we first looked at its subcellular distribution. As shown recently for other tissues (Liu et al., 2017; Raote et al., 2017), Tango1 assembles into ring-like structures in tracheal terminal cells, and co-localizes with the ERES marker Sec16 (Fig. 1E). The truncated Tango1^2L3443^ protein fails to colocalize with Sec16, and Sec16 distribution itself is also altered in *tango1^2L3443^* mutant cells and upon *tango1* knockdown (Fig. 1F and Fig. S1E). While in control cells Sec16 particles show a homogenous distribution with a narrow range of sizes with a mean/median of 0.54μm^2^/0.49μm^2^, cells lacking Tango1 contain larger range of sizes with a mean/median of 0.44μm^2^/0.29μm^2^ (Fig. 1E-F, S1D-E).

Golgi morphology is also abnormal in *tango1^2L3443^* cells, as shown by the distribution of the Golgi marker ManII-GFP relative to Sec16. In control cells, Sec16 and ManII-GFP are seen as juxtaposed spots, whereas in *tango1^2L3443^* mutant cells ManII-GFP seems to enclose Sec16 particles (Fig. S1F-G), consistent with previous studies suggesting the retention of ManII-GFP near the ER (Bard et al., 2006). RNAi against *tango1* in terminal cells also induced abnormal aggregation of the Golgi marker ERGIC53 (Fig. 1G-H).

### The role of Tango1 in terminal cells

Tango1 has been studied for its role in the trafficking of collagen in cultured mammalian cells and in Drosophila fat body cells, the main collagen producers in the fly (Pastor-Pareja and Xu, 2011; Saito et al., 2009). Terminal cells are surrounded by collagen, and although according to expression data collagen may be expressed only at minimal levels in tracheal cells, it was possible that the defects seen in tracheal cells might be due to failures in the secretion of collagen. To test this, we knocked down collagen (encoded by the gene *viking, vkg*) specifically in terminal cells. This did not result in any morphological defects of the type that loss of Tango1 caused (Fig. S2A). We also compared the effects of knocking down *tango1* either in terminal cells or in the fat body. We found that collagen levels surrounding terminal cells are affected only when *tango1* is knocked down in the fat body but not when it is absent in terminal cells (Fig. S2B-G). These experiments show first that the collagen surrounding terminal cells is not produced by the terminal cells but mostly, if not entirely, by the fat body, and secondly, that the defects resulting from *tango1* loss-of-function in terminal cells cannot be explained by a defect in the transport of collagen.

If the defects in *tango1* mutant terminal cells cannot be explained by failure of collagen secretion, then they must be due either to a failure to transport to the cell surface other molecules essential for tracheal function, or to a function unrelated to the secretion of specific substrates (for example, a global failure within the secretory pathway).

We analysed the distribution of a range of cell surface and secreted proteins in Tango1-depleted terminal cells. These included markers for the basal and apical cell membranes, because morphological defects in epithelial cells are often associated with defective cell polarity. The localization of bPS integrin at the outer, basal membrane of the cell was not affected by *tango1* knockdown (Fig. 2F-G). By contrast, the apical membrane protein Crumbs (Crb), normally present at the luminal plasma membrane (Fig. 2H), failed to reach its normal destination and was instead found dispersed throughout the cytoplasm (Fig. 2I). These observations favour a role for Tango1 in the transport of specific proteins rather than in general secretion.

Since the suggested role for Tango1 is to aid the secretion of very large cargos, we examined the distribution of Dumpy (Dpy), the largest protein encoded in the Drosophila genome, with a size of 2.5 MDa and a length of 800 nm (Misra et al., 2002; Wilkin et al., 2000). Dpy contains EGF-repeat domains and a Zona Pellucida (ZP) domain. It mediates the attachment between cells and the chitinous apical extracellular matrix (aECM), through its interaction with Pio, a ZP transmembrane protein (Ozturk-Colak et al., 2016). We visualized Dpy through a YFP insertion at the *dpy* locus that results in a fusion protein expressed at endogenous levels, Dpy-YFP (Ray et al., 2015).

In cells of the tracheal dorsal trunks we distinguished two pools of Dpy: one that was secreted and was seen within the lumen of the trachea, the other in the cytoplasm, in the form of spots, which were presumably vesicles containing Dpy on its secretion route (Fig. 2J). We found that a subset of the Dpy particles was partly or fully surrounded by Tango1 and in close proximity to the Golgi marker GM130 (Fig. 2J, insets). In terminal cells, Dpy-YFP is present in the lumen of the cells, where it is enriched at the plasma membrane, together with its binding partner Pio (Fig. 2K). In *tango1* knockdown terminal cells, neither Dpy-YFP nor Pio were found in the lumen, and they instead accumulated in the cytoplasm (Fig. 2L).

To test whether the mislocalisation of any of the molecules that we analysed was responsible for the defects seen in tracheal cells, we depleted Dpy and Pio from terminal cells. Neither *dpy* nor *pio* knockdown produced air-filling defects or a reduction in the number of branches (Fig. 2C-E), in spite of efficient silencing of *pio* expression (Fig. S3A-B). Therefore the morphological defects resulting from loss of Tango1 cannot be explained by inefficient Dpy or Pio secretion. Similarly, mislocalisation of Crb is not sufficient to explain the *tango1* loss-of-function phenotype, since *crb* homozygous mutant terminal cells do not show branching defects that resemble the *tango1* phenotype (Schottenfeld-Roames et al., 2014).

In summary, regarding the dependence of different cargos on Tango1 we have found three cases: Dpy represents a cargo that fits the expected characteristic of Tango1 substrates of being very large; Crb is a cargo that depends on Tango1 although it is not large; and finally, bPS integrin is a cargo that does not depend on Tango1. To learn more about the rules and generalities of Tango1-dependent secretory cargos, we examined other tissues.

### Effect of Tango1 loss-of-function on Dpy in wing discs, glial cells and tendons

Dpy serves as a scaffold that anchors tissues to the aECM and supports tissue shape changes in many organs, and its function has been most extensively studied in the wing disc (Ray et al., 2015). Knocking down *tango1* in the wing pouch resulted in intracellular accumulation of Dpy-YFP (Fig. 3A, E). Loss of Tango1 was again associated with changes in the distribution of Sec16. This was particularly evident when *tango1* was knocked down in a stripe across the disc using *ptc-gal4*. We found that in the absence of Tango1, the number of Sec16 particles per area was reduced (Fig. 3B, F).

**Figure 3.**
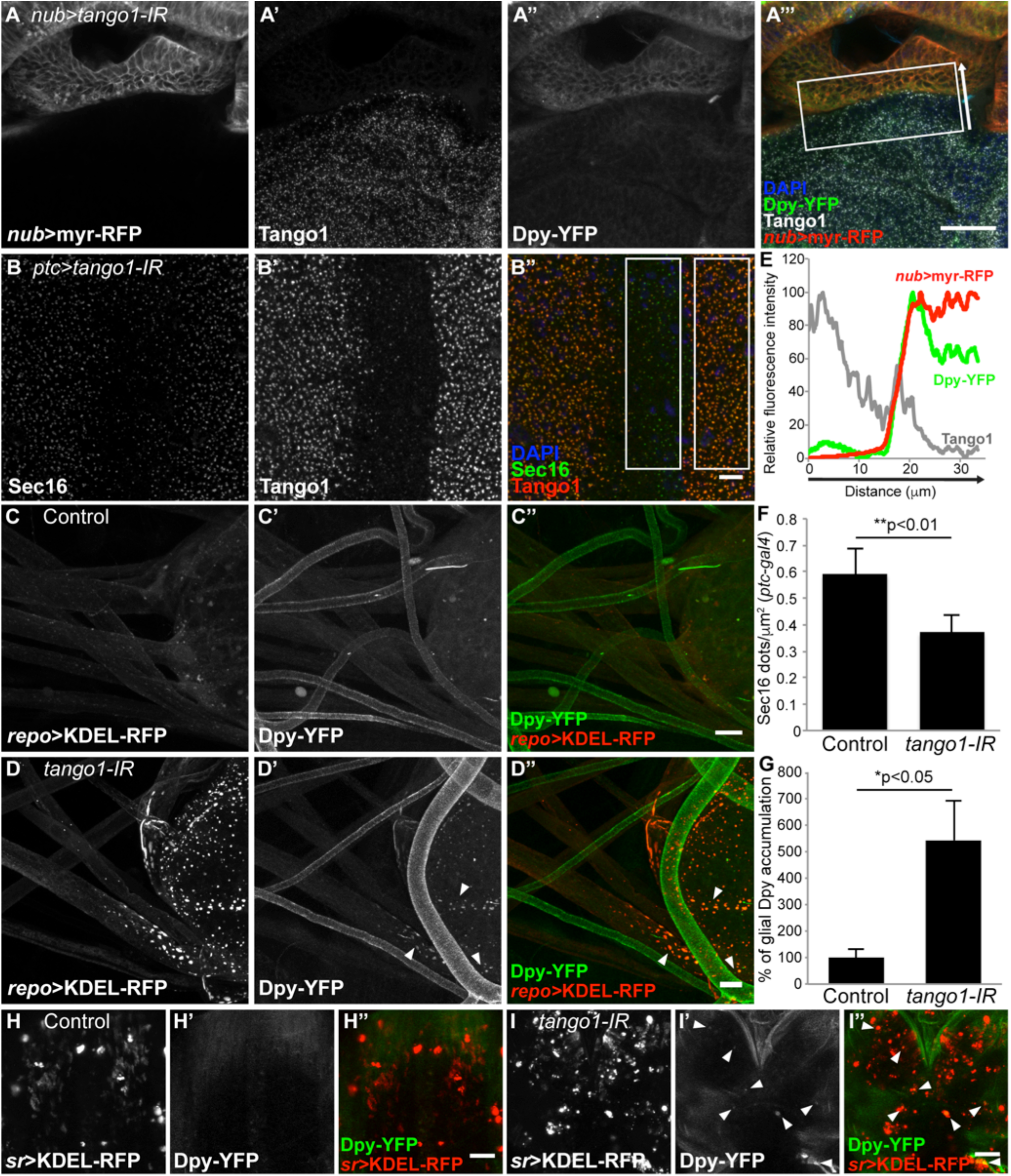
Cargo accumulation and ER defects in wing disc, glial and tendon cells. (A) Wing disc from animals with an endogenously tagged Dpy protein (Dpy-YFP) expressing myr-RFP and *tango1-IR* in the wing pouch under *nub-gal4*. Tango1 was stained to confirm the efficiency of knockdown. (B) Wing disc expressing *tango1-IR* under the *ptc-gal4* driver and stained for Sec16. Absence of Tango1 staining reveals the region where *tango1-IR* is expressed. (C, D) Larval brains expressing KDEL-RFP as an ER marker under the glial-specific driver *repo-gal4*. Arrowheads in (D) show sites of Dpy accumulation, and their quantification is shown in (G). (E) Intensity profile in the direction of the white arrow and summed across the width of the box in (A). (F) The number of Sec16 dots (+/− SD) in the boxed regions in (B’’). Number of discs analyzed = 4. Significance was determined using two-tailed t-test. (G) Quantification of the level of Dpy-YFP (+/− SD) retained within the repo>KDEL-RFP channel. Control, n=4; *tango1-IR*, n=4. Significance was determined using two-tailed t-test. (H, I) Live pupae expressing KDEL-RFP under the tendon cell driver *sr-gal4*, and endogenous Dpy-YFP expression. Arrowheads in (I) point to sites of Dpy accumulation. Scale bars are 25μm (A), 5μm (B), 10μm (C, D), and 50μm (H, I).

Tango1 has previously been shown to be active in larval glial cells and pupal tendons (Petley-Ragan et al., 2016; Tiwari et al., 2015) and we found that these cell types are surrounded by Dpy-YFP (Fig. 3C, H), consistent with expression reports on other developmental stages (Knowles-Barley et al., 2010; Wilkin et al., 2000). Depletion of Tango1 resulted in the intracellular accumulation of Dpy-YFP in both tissues (Fig. 3D, G, I). We compared the localization of the intracellular Dpy-YFP spots in glial and tendon cells with that of KDEL-RFP, an ER marker. Dpy-YFP co-localized with KDEL-RFP in cells lacking Tango1, suggesting Dpy remains within the ER in these cells (Fig. 3C-D, H-I). These experiments indicate that the role of Tango1 in Dpy secretion is general, and not restricted to tracheal cells. Whereas the tissues studied so far each have their own, specific cargos that depend on Tango1, they also share Dpy as a common cargo.

### Direct and indirect effects of loss of Tango1 on cargo accumulation in the fat body

Tango1-dependent trafficking has been most thoroughly characterized in the fat body. In fat body cells lacking Tango1, a number of cargos including collagen are not delivered to the cell surface, and the structure of the ER and Golgi are abnormal (Liu et al., 2017; Pastor-Pareja and Xu, 2011). Fat body cells do not express Dpy, and as in tracheal cells, endogenous bPS integrin distribution is not affected by lack of Tango1 (Fig. 4A-B).

**Figure 4.**
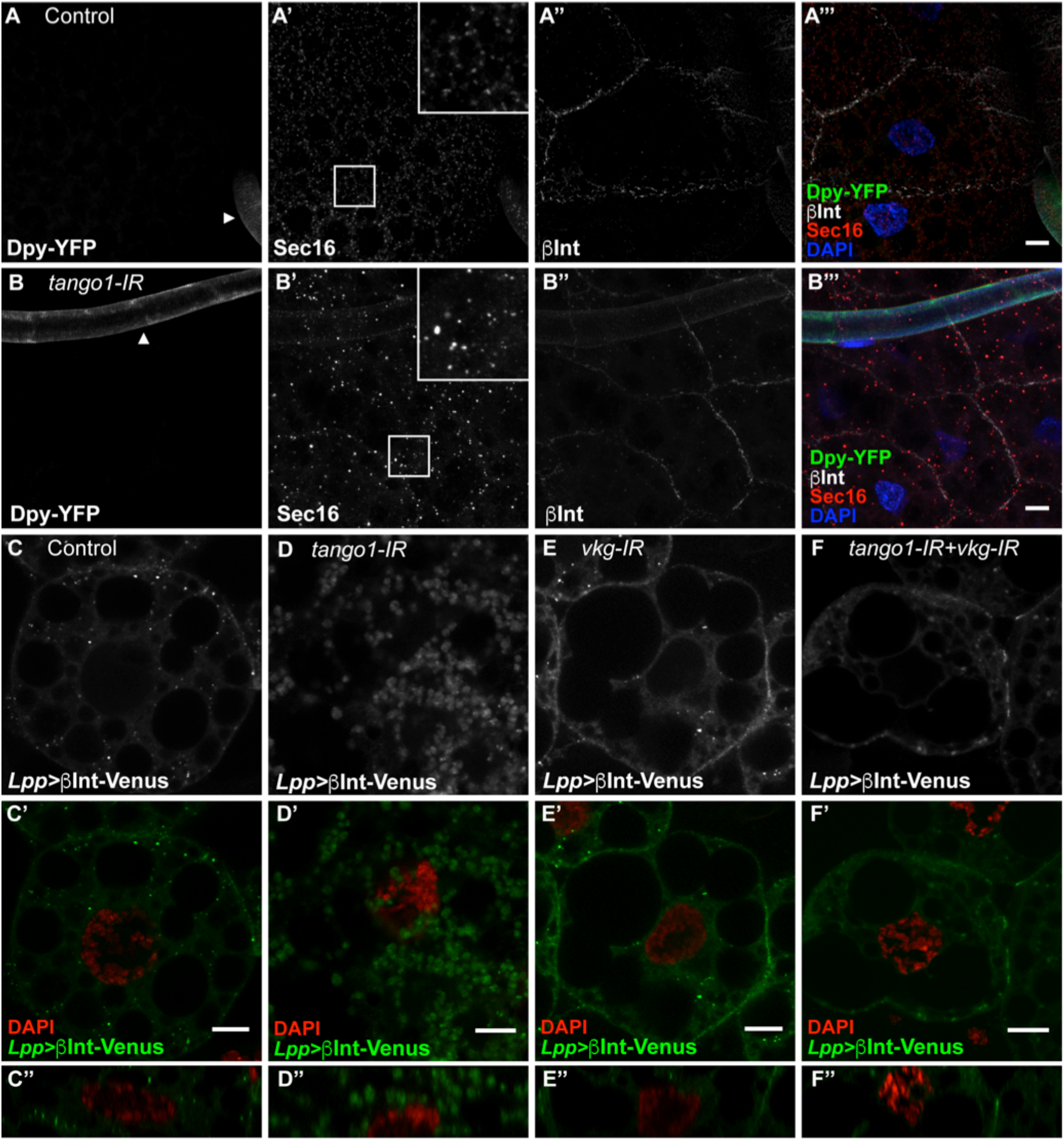
Effect of loss of *tango1* in the fat body on βPS integrin and collagen. (A-B) Single z sections of fat body cells from Dpy-YFP larvae were stained for Sec16 and βPS integrin (βInt). Arrowheads point to tracheal tubes (not affected by transgenes expressed under *Lpp-gal4*) as positive control for Dpy-YFP expression. In the absence of *tango1* (B), the regular distribution of Sec16 is lost, whereas βInt is not affected. (C-F) Single z sections of fat body cells expressing βInt-Venus under *Lpp-gal4*. Control cells (C) are able to deliver βInt-Venus to the cell membrane, whereas *tango1-IR* cells (D) cannot. The absence of collagen (*vkg-IR*, E) does not affect βInt-Venus delivery. (F) Knocking down both *tango1* and *vkg* rescues membrane delivery of βInt-Venus. (C’’-F’’) Orthogonal views of the same cells. Scale bars are 10μm.

We noticed that independent of size, secretion of several overexpressed molecules was impaired upon *tango1* knockdown in fat body cells. This included Gasp-GFP, with a molecular weight of only 55 kDa (Fig. S3C-D), and overexpressed βPS integrin-Venus, even though endogenous βPS integrin was unaffected (Fig. 4C-D). Previous reports have also shown that in Drosophila, the absence of Tango1 leads to the accumulation of other overexpressed small cargos like secreted HRP and GFP, and of Hedgehog-GFP (Bard et al., 2006; Liu et al., 2017). This was also observed in cultured mammalian cells for secreted HRP and GFP (Nogueira et al., 2014; Saito et al., 2009). In the case of mammalian cells, it was suggested that HRP accumulation was caused by unsecreted collagen blocking the secretory pathway (Nogueira et al., 2014). To test whether such a mechanism may explain the failure of smaller molecules to be secreted in *tango1*-deficient fat body cells, we studied whether the reduction of *vkg* would improve the secretion of small cargos by simultaneously knocking down *vkg* and *tango1*. We found that if in addition to *tango1*, we knocked down *vkg*, this resulted in the rescue of the secretion of overexpressed βPS integrin (Fig. 4E-F) as well as overexpressed Crb fused to GFP (Crb-GFP, Fig. S4A-D). To exclude an artefactual amelioration of the *tango1*-knockdown phenotype because the second RNAi construct might reduce the efficiency of *tango1* knockdown, in this and further experiments we compared Tango1 and collagen levels in the double knockdown condition with individual *tango1* and *vkg* knockdowns and found that both targets were equally well silenced in the two conditions (Fig. S4E-H). These results show that both overexpressed βPS integrin-Venus and Crb-GFP can be delivered to the membrane in the absence of Tango1 if collagen is also removed, suggesting that their accumulation upon *tango1* knockdown is an indirect effect of collagen accumulation.

We also wanted to test whether SPARC and laminins, two other cargos known to depend on Tango1 for their secretion [(Petley-Ragan et al., 2016; Tiwari et al., 2015), Fig. 5A-B, E-F and Fig. S3E-F] might be blocked by collagen accumulation in the fat body. These experiments were inconclusive because loss of collagen itself lead to laminins and SPARC retention in the ER (Fig. 5C, G; Fig. S3G), and simultaneous collagen and Tango1 knockdown therefore did not rescue laminin or SPARC secretion (Fig. 5D, H; Fig. S3H).

**Figure 5.**
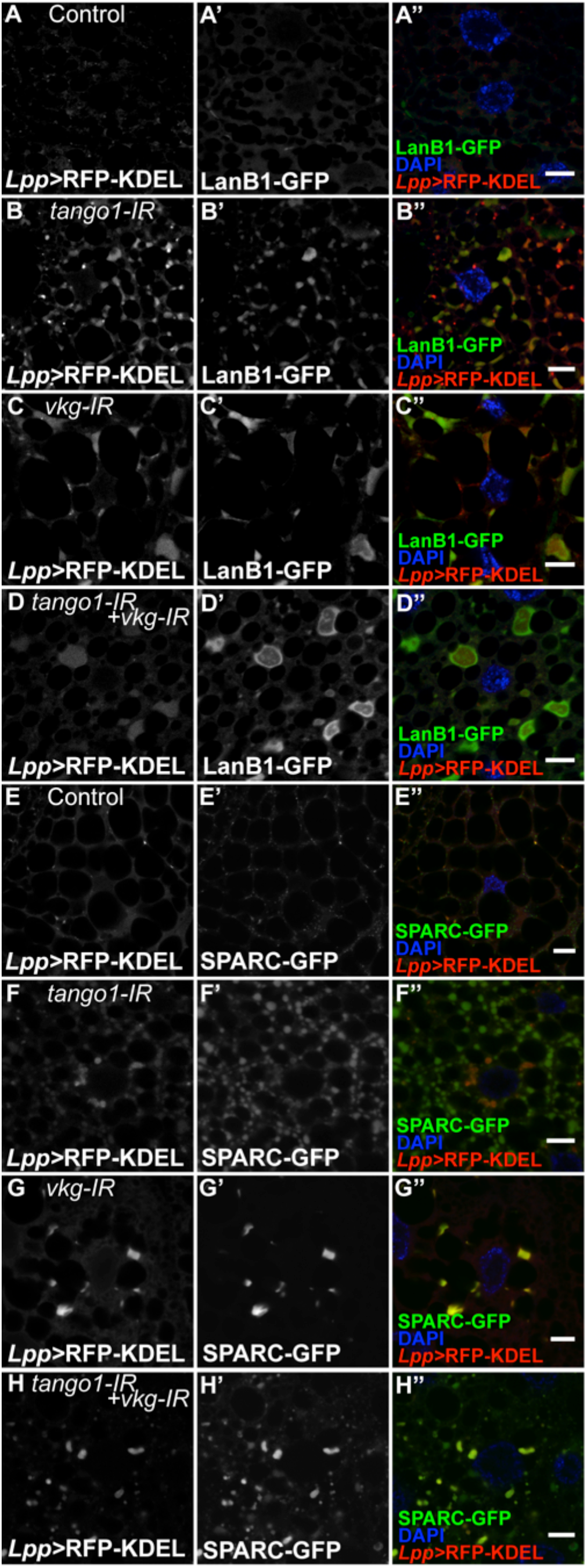
Dependence on collagen of Laminin and SPARC secretion in fat body cells. KDEL-RFP was expressed in fat body cells using *Lpp-gal4* in animals expressing LanB1-GFP (A-D) or SPARC-GFP (E-H) under their endogenous promoters (fTRG library). Both proteins are retained in the ER in the absence of Tango1 (B, F), collagen (C, G) or both (D, H). Scale bars are 10μm.

### Direct and indirect effects of loss of Tango1 on cargo accumulation in glial cells and in terminal cells

Like overexpressed βPS integrin and Crb in the fat body, some of the Tango1-dependent cargos identified in tracheal, glial, wing epithelial and tendon cells are also not particularly bulky. We therefore investigated whether they might also not be direct substrates of Tango1. These tissues do not express detectable levels of collagen (Pastor-Pareja and Xu, 2011; Petley-Ragan et al., 2016; Tiwari et al., 2015), and it was therefore unlikely that unsecreted collagen was the blocking cargo. We therefore wondered whether Dpy, as another large Tango1 cargo might be blocking the secretory pathway.

Glial cells of the larval brain and of the peripheral nervous system have also been shown to need Tango1 for the secretion of laminin chains LanB1 and LanB2 (Petley-Ragan et al., 2016). Laminins are assembled into trimers composed of the LanB1, LanB2 and LanA subunits. All subunits are required for trimer secretion, but LanA can also be secreted as a monomer. We found that as has been shown for LanB1 (Fig. S3I-J), *tango1* knockdown also resulted in LanA accumulation in the ER (Fig. 6A-B, E). It was puzzling that LanA was retained at the ER in glial cells lacking Tango1, considering that it should be able to be secreted as a monomer even when LanB1 and LanB2 are not secreted (Hamill et al., 2009). To test if accumulated intracellular Dpy might be responsible for this, we knocked down *tango1* and *dpy* simultaneously. We found that the defective secretion of both LanA and LanB1 caused by lack of Tango1 was rescued by also silencing *dpy* (Fig. 6D-E, Fig. S3L; controls for knockdown efficiency in Fig. S5A-D).

**Figure 6.**
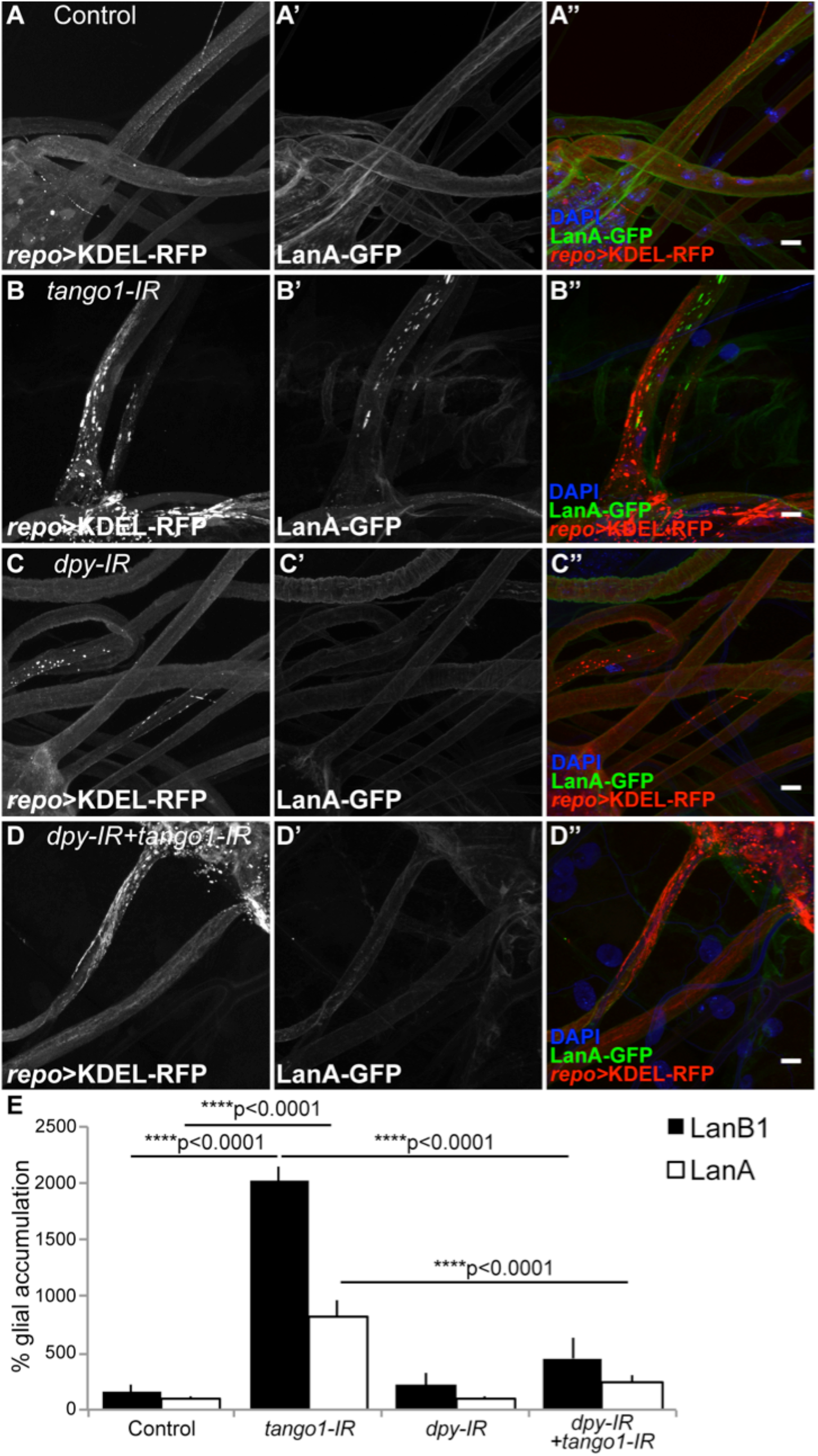
Laminin accumulation in *tango1*-depleted cells can be suppressed by removing Dpy. KDEL-RFP was expressed in glial cells using *repo-gal4* in animals expressing LanA-GFP under its endogenous promoter (fTRG library) in control cells (A), in cells expressing *tango1-IR* (B), *dpy-IR* (C) or both (D). *tango1-IR* induces LanA-GFP retention at the ER (B). While *dpy-IR* alone does not affect LanA-GFP distribution (C), it suppresses the *tango1-induced* LanA-GFP accumulation (D). (E) Quantification of the intracellular level of LanA and LanB1 with respect to that of control animals +/− SEM. Control, LanA n=4, LanB1 n=3; *tango1-IR*, LanA n=5, LanB1 n=3; *dpy-IR*, LanA n=6, LanB1 n=3; *tango1-IR+dpy-IR*, LanA n=7, LanB1 n=3. Significance was determined using one-way ANOVA and Tukey’s multiple comparisons test. Scale bars are 10μm

In tracheal cells, where Crb delivery to the membrane was completely abolished by *tango1* knockdown (Fig. 2H-I, Fig. 7A-B), the additional knockdown of *dpy* caused Crb membrane localization to be re-established (Fig. 7D, controls for knockdown efficiency in Fig. S5E-H). In summary, the effect of loss of Tango1 on a broad range of cargos is indirect, with the proximal effect being the retention of one or perhaps a small number of direct substrates, which in turn blocks the proper trafficking of other molecules.

**Figure 7.**
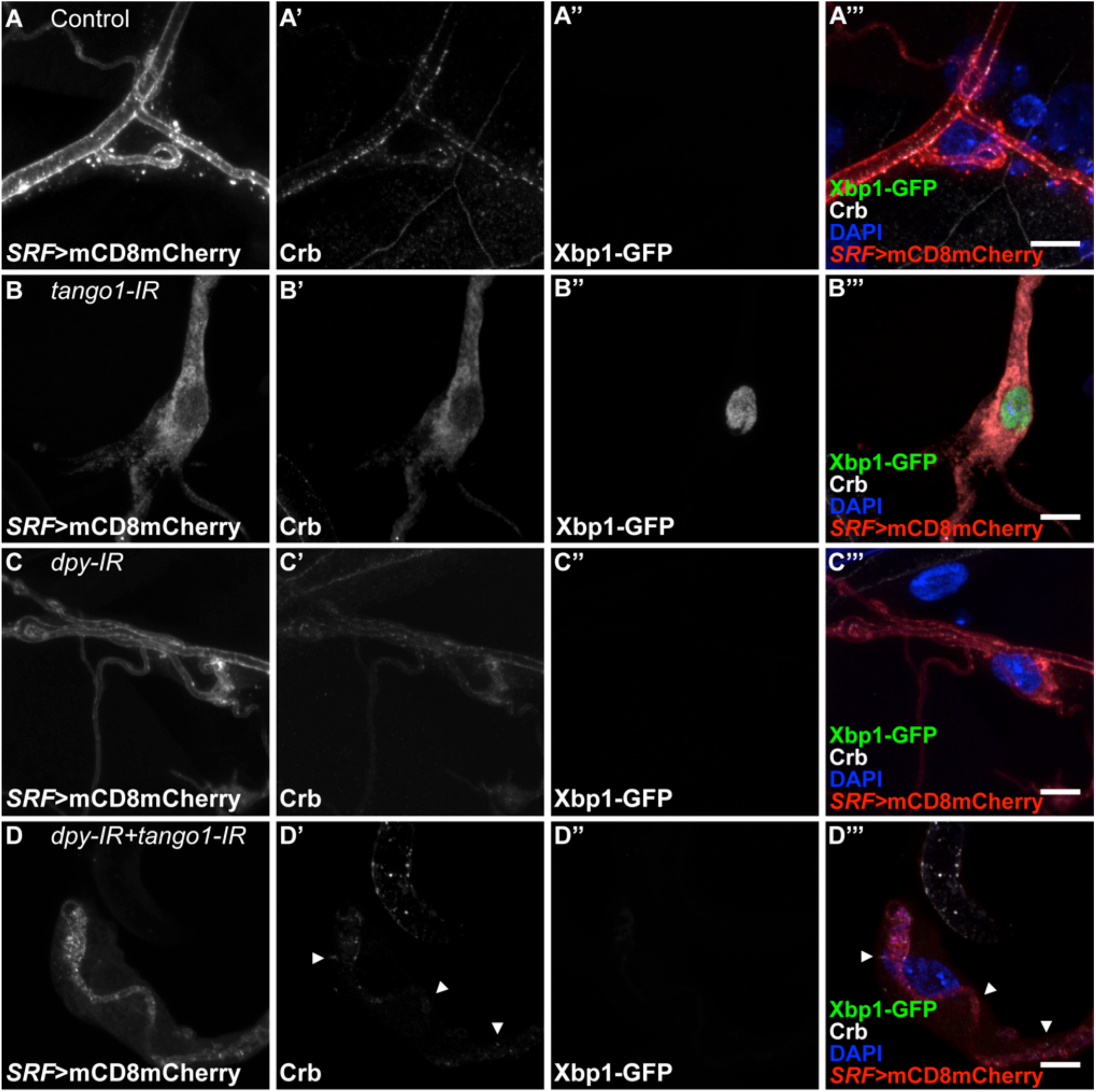
Effect of *tango1* and *dpy* knockdown on Crumbs and the ER stress response. Terminal cells expressing mCD8mCherry and Xbp1-GFP under *SRF-gal4*. Xbp-1-GFP is translated and accumulated in the nucleus only after activation of the ER-stress response (Ryoo, et al., 2007). In control (A) and *dpy-IR* cells (C), Crb localizes to the luminal membrane and Xbp1-GFP is not detectable. In *tango1-IR* cells (B), Crb is not able to localize to the luminal membrane and Xbp1-GFP accumulates in the nucleus. These defects can be suppressed by additionally knocking down *dpy* (D). Scale bars are 10μm.

### Tango1 loss-of-function in terminal cells leads to ER stress in a Dpy-dependent manner

Our results so far pointed towards *tango1* loss-of-function in terminal cells and in other tissues affecting not only the secretion of its own cargo, but also that of others as a side effect of the ER being clogged by unsecreted cargo. Another documented consequence of loss of Tango1 is the activation of the ER stress response (Petley-Ragan et al., 2016). One of the inducers of ER stress is protein retention in the ER, and we therefore investigated whether the ER stress response was a direct consequence of loss of Tango1, or instead, the result of abnormal protein accumulation in the ER. To test this, we used the marker Xbp1-GFP, which is post-transcriptionally upregulated in response to ER stress (Coelho et al., 2013; Ryoo et al., 2007). We observed high levels of Xbp1-GFP in terminal cells lacking Tango1, but not in control or *dpy-IR* cells (Fig. 7A-C). When *tango1* and *dpy* were simultaneously depleted, Xbp1-GFP was no longer upregulated (Fig. 7D). Therefore, the ER stress response is also not a direct consequence of loss of Tango1, but the result of the incorrect trafficking of Dpy.

The original phenotype for which we identified the *tango1* mutation was defective terminal cell morphology. We argued above that this was not due to the loss of Dpy or Pio at the cell surface (Fig. 2C-E). Having found that ER stress and indirect retention of small cargos could be suppressed by removing primary cargos, we wondered whether the morphological defects were also secondary to protein accumulation, or if they revealed a direct function of Tango1 independent of its role in secretion of Dpy. We found that the defects of branch number and branch air filling seen in *tango1* knockdown cells were significantly suppressed if *dpy* was also silenced (Fig. 8). This suggests that most of the deleterious effect of Tango1 depletion on terminal cell morphology is a consequence of abnormal Dpy accumulation. Since loss of *dpy* itself has no effect on cell morphology, Dpy and the resulting failure in protein secretion or ER stress may be the cause for the defects in cell shape.

**Figure 8.**
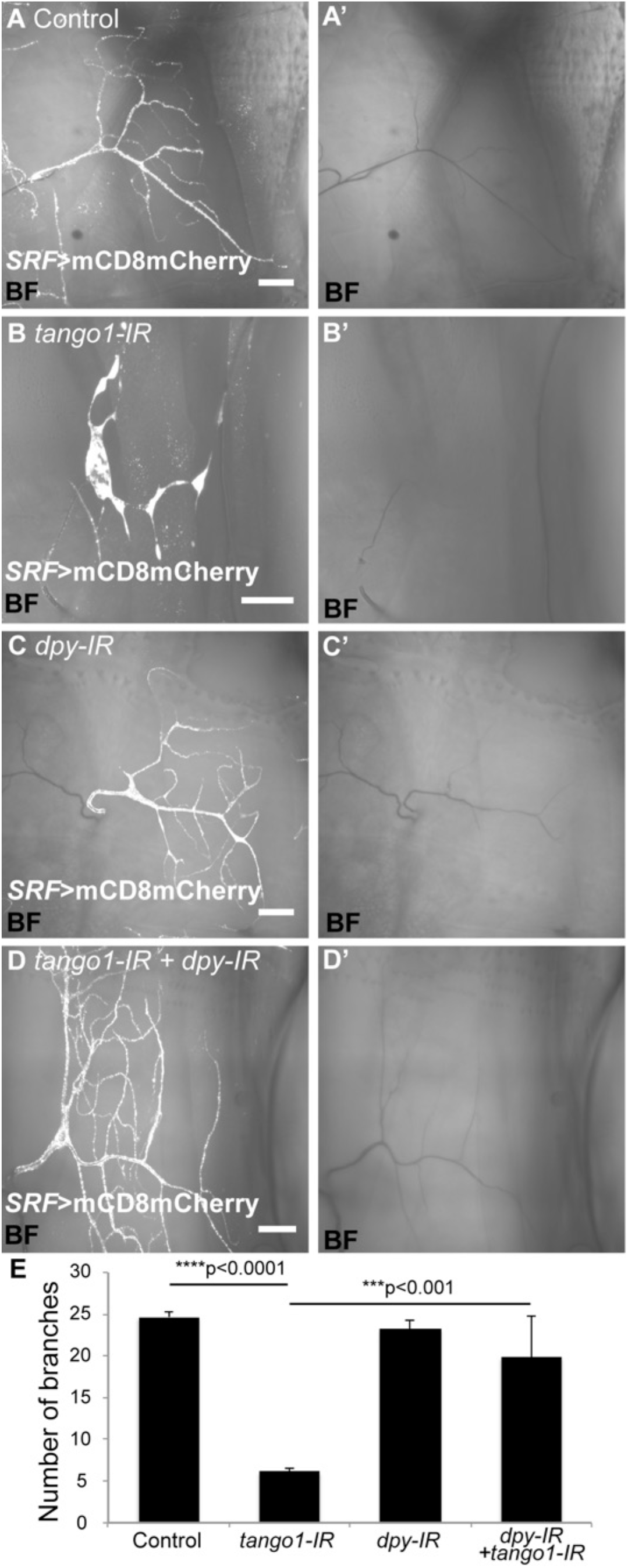
Rescue of cell morphology in *tango*–depleted cells by removal of Dpy. (A-D) Bright field (BF) and mCD8mCherry expression under the terminal cell-specific driver *SRF-gal4*. In control (A) and *dpy-IR* cells (C), branches are filled with gas, whereas the absence of *tango1* leads to failure of air-filling and reduced branching (B). Both defects are suppressed by additionally knocking down *dpy* (D). (E) Quantification of branching in (A-E). Bars represent mean +/-SEM. Control, n=4; *tango1-IR*, n=9; *dpy-IR* n=8; *tango1-IR+dpy-IR*, n=8. Significance was determined using one-way ANOVA and Tukey’s multiple comparisons test.

### Separable roles for Tango1 on ER/Golgi architecture and secretion

A further defect others and we had observed to result from loss of Tango1 was the disruption of the normal organization of the Golgi and ER. Since the other defects we described above – failure to secrete a range of proteins and defective cell morphology – were indirect effects of Tango1 we wondered whether this might also be true for the distorted Golgi and ER, and whether these might also be caused by large cargo accumulation (in this case Dpy). However, we found that upon double *tango1* and *dpy* RNAi, Sec16 organization was not restored and we still observed a wide range of Sec16 particle sizes and staining intensities (Fig. 9D-F). Similar results were obtained for GM130, a Golgi marker; while control and *dpy-IR* cells showed a homogeneous size distribution of GM130-stained particles (Fig. 10A, C), in *tango1* and double knockdown cells the distribution of GM130 is altered (Fig. 10B, D). Given that in the double knockdown cells Dpy is no longer clogging the ER and that secretion of other molecules is re-established, these results indicate that Tango1 has an additional function in maintaining ER-Golgi morphology that is independent of its role in bulky cargo transport.

**Figure 9.**
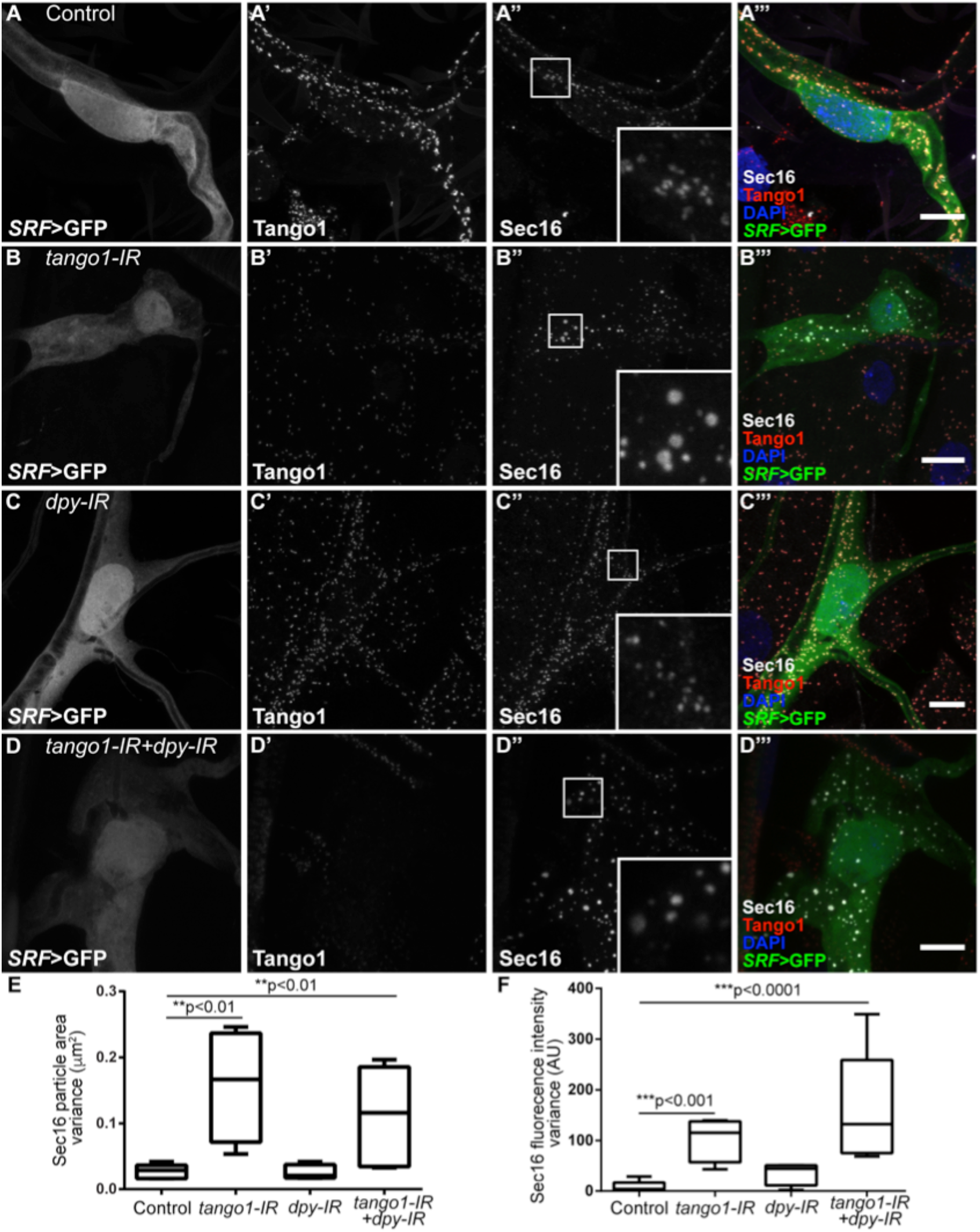
Effect of loss of Tango1 and Dpy on Sec16 distribution. (A-D) Sec16 in terminal cells expressing GFP under *SRF-gal4*. In control (A) and *dpy-IR* cells (C), Sec16 particles are homogeneous in size and fluorescence intensity; in *tango1-IR* cells Sec16 particle size and fluorescence intensity is variable (B) and this variability is not altered by simultaneously removing Dpy (D). (E-F) Variance of Sec16 particle size (E) and of Sec16 fluorescence intensities (F). Control, n=5; *tango1-IR*, n=4; *dpy-IR*, n=4; *tango1-IR+dpy-IR*, n=5. Significance was determined using one-way ANOVA and Sidak’s multiple comparisons test. Scale bars are 10μm.

**Figure 10.**
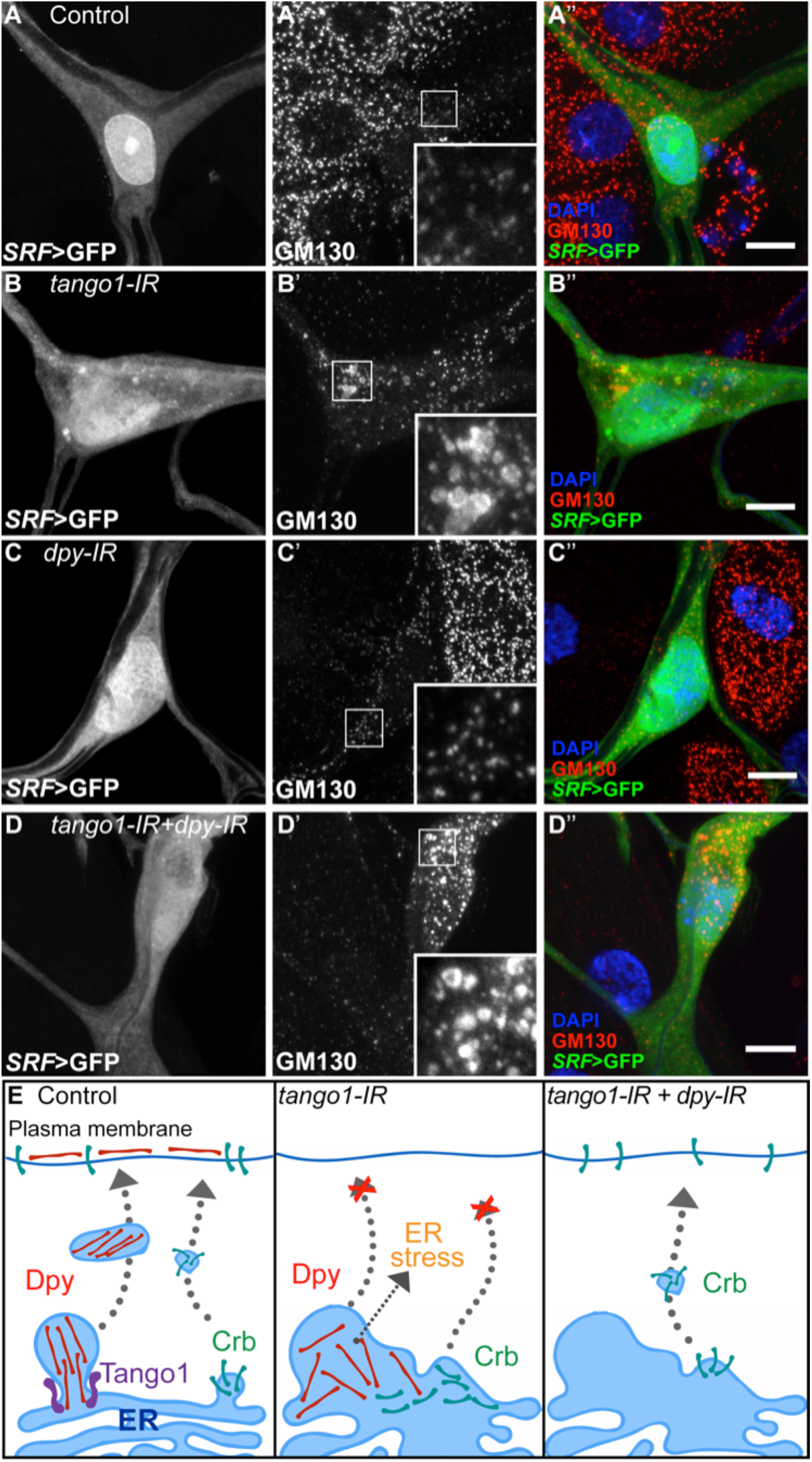
Effect of loss of Tango1 and Dpy on GM130 distribution. (A-D) Terminal cells expressing GFP under *SRF-gal4*, and stained for the Golgi marker GM130. In control (A) and *dpy-IR* cells (C), the distribution and size of GM130-labelled structures is homogeneous whereas in *tango1-IR* cells GM130 is seen in heterogeneous aggregates (B). Knocking down *dpy* in tango1-IR cells does not rescue GM130 distribution (D). Scale bars are 10μm. (E) Model showing the role of Tango1 in Dpy trafficking and the indirect consequences of Dpy blockage. In the absence of Tango1, the structure of the ER is changed, the ER stress response is activated and neither Dpy nor Crb reach the plasma membrane. If Dpy levels are reduced, the ER stress response is no longer active and Crb can be secreted. However, the ER morphology is not restored.

## Discussion

We have described a novel role of Tango1, which we initially identified through its function in tracheal terminal cells and other tissues in Drosophila embryos, larvae and pupae. Due to their complex shapes and great size, terminal cells are a well suited system to study polarized membrane and protein trafficking, with the easily scorable changes in branch number and maturation status providing a useful quantitative readout that serves as a proxy for functional membrane and protein trafficking machinery. Moreover, our analyses are conducted in the physiological context of different tissues in the intact organism.

### Nature of the *tango1^2L3443^* allele

The loss of function allele *tango1^2L3443^* has a stop codon 8 amino acids downstream of the PRD domain, and eliminates the 89 C-terminal amino acids of the full-length protein. It is unlikely that the mutation leads to a complete loss of function. First, terminal cells expressing an RNAi construct against *tango1* show stronger defects, with fewer branches per cell than homozygous *tango1^2L3443^* cells. Secondly, the mutant protein appears not to be destabilized nor degraded, but instead is present at apparently normal levels, albeit at inappropriate sites. Predictions of the deleted fragment of the protein suggest it is disorganized and that it contains an arginine-rich domain that has no known interaction partners and that is not present in human Tango1. In homozygous mutant terminal cells the mutant Tango1^2L3443^ protein fails to localize at ERES. In mammalian Tango1, the PRD domain is necessary for the localization of Tango1 to the ERES and for its interaction with Sec23 and Sec16 (Maeda et al., 2017; Saito et al., 2009), but since this domain is fully present in Tango1^2L3443^, our results mean that either the missing 89 C-terminal amino acids contain additional essential localization signals, or that the PRD domain is structurally affected by the truncation of the protein. We consider the latter less likely, as a truncation 8 amino acids downstream of the PRD domain is unlikely to destabilize the poly-proline motifs, especially as the overall stability of the protein does not seem to be affected. Furthermore, this region shows a high density of phospho-serines [Ser-1345, Ser-1348, Ser-1390 and Ser-1392 (Zhai et al., 2008)] suggesting it might serve as a docking site for adapter proteins or other interactors.

### Possible causes of the cellular morphological defects

Terminal cells lacking Tango1 have fewer branches than control cells, and are often not properly filled with air. This loss-of-function phenotype is not due to a direct requirement for Tango1, as it is suppressed by the simultaneous removal of Dpy. It also cannot be explained by the individual loss of *crb, pio or dpy*, since knocking down any of these genes has no effect on cell morphology. It is possible that the combined loss of these, and perhaps further proteins we have not tested, at the apical membrane might lead to defective branch formation or stability, but we believe the phenotype is more likely a secondary consequence of the general defects caused by loss of Tango1 and the accumulation of Dpy in the ER. For example, these defects might lead to a failure to deliver sufficient lipids and membrane from the ER to the apical plasma membrane. Alternatively, the activation of the ER stress response that we observe upon loss of *tango1* might have additional side effects on cell morphology.

### Dumpy, a new cargo of Tango1

Collagen, with a length of 300 nm and ApoB chylomicrons with a diameter of >250 nm, have both been biochemically validated as Tango1 cargos (Saito et al., 2009; Santos et al., 2016). These molecules are not expressed in terminal cells [this work and (Baer et al., 2012)], and therefore it was clear that Tango1 must have a different substrate in these cells. Given that Tango1 is known for the transport of bulky cargo, that Dpy is the largest Drosophila protein at 800 nm length, and that Dpy vesicles are associated with Tango1 rings in tracheal cells, we propose that Dpy is a further direct target of Tango1. Colocalization of Tango1 with its cargo has also been observed in other tissues: with collagen in Drosophila follicle cells and with ApoB in mammalian cell lines (Lerner et al., 2013; Santos et al., 2016). Proteomic studies have shown an indirect interaction between Dpy and Sec16, supporting a COPII-mediated transport of Dpy (Rees et al., 2011).

No regions of sequence similarity that could represent Tango1 binding sites have been found in Tango1 cargos. There are several possible explanations for this. First, these proteins may contain binding motifs, but the motifs are purely conformational and not represented in a linear amino acid sequence. There is no evidence for or against this hypothesis, but it would be highly unusual, and there is support for alternative explanations. Thus, as a second possibility, all three proteins may require Tango1 for their secretion, but variable adapters could mediate the interactions. In vertebrates, Tango1 can indeed interact with its cargo through other molecules; for instance, its interaction with collagen is mediated by Hsp47 (Ishikawa et al., 2016). However, in Drosophila there is no Hsp47 homolog (Martinek et al., 2008). In the case of ApoB, it has been suggested that microsomal triglyceride transfer protein (MTP) and its binding partner, protein disulphide isomerase (PDI), might associate with Tango1 and TALI to promote ApoB chylomicrons loading into COPII vesicles. Evidence supporting this is that the lack of MTP leads to ApoB accumulation at the ER (Pfeffer, 2016; Santos et al., 2016). It is not known if secretion of other Tango1 cargos like collagen or Dpy also depends on MTP and PDI, but PDI is known also to form a complex with the collagen-modifying enzyme prolyl 4-hydroxylase (Kivirikko and Myllyharju, 1998). We have previously shown that terminal cells lacking MTP show air-filling defects and fail to secrete Pio and Uninflatable to the apical membrane, and that loss of MTP in fat body cells also affects lipoprotein secretion (Baer et al., 2012), as it does in vertebrates. Since cells lacking MTP or Tango1 have similar phenotypes, it is plausible that the MTP function might be connected to the activity of Tango1.

### Clogging of the ER

We interpret our data to mean that in the absence of Tango1, primary cargo accumulates in the ER, and in addition, there are secondary, indirect effects that can be suppressed by reducing the Tango1 cargo that overloads the ER. The secondary effects include activation of the ER stress response and intracellular accumulation of other trafficked proteins like Crb, laminins, and overexpressed proteins and probably also the accumulation of heterologous proteins like secreted HRP or GFP in other systems (Nogueira et al., 2014).

We can think of two explanations for how accumulation of Tango1 cargo might affect the secretion of other proteins. Primary cargo accumulating in the ER could be inhibiting the secretion of other proteins by blocking access of all proteins in the ER to ERES. However, we find this unlikely given that βPS integrin can still be secreted towards the plasma membrane in these cells. An alternative explanation may be an involvement of the ER stress response. While the ER stress response does not seem to be a direct consequence of loss of Tango1 since it can be suppressed by removing ER overload, the activation of the ER stress response might nevertheless actively affect the secretion of certain proteins from the ER to the Golgi apparatus.

### Different sensitivities of bPS integrin and Crb to loss of Tango1

It is not immediately clear why cargo accumulation in terminal cells lacking Tango1 affects the secretion of Crb but not of βPS integrin. While we look at steady states in our analyses, Maeda et al. have measured the dynamics of secretion and find that loss of Tango1 leads to a reduced rate of secretion of VSVG-GFP, an effect that we would have missed for any proteins we classify as not affected by loss of Tango1 (Maeda et al., 2017). Irrespective, we can think of a range of mechanisms that might be responsible for this difference, including alternative secretion pathways and differences in protein recycling. Alternative independent secretory pathways have been reported in different contexts. For instance, while both αPS1 and βPS integrin chains depend on Sec16 for their transport, the αPS1 chain can bypass the Golgi apparatus and can instead use the dGRASP-dependent pathway for its transport (Schotman et al., 2008). It would be possible then that in terminal cells, βPS integrin is also trafficked through an alternative pathway that is not affected by loss of Tango1. Similarly, tracheal cells lacking Sec24-CD (encoded by the gene *gho*) accumulate Gasp, Vermiform and Fasciclin III, but not Crb (Norum et al., 2010), supporting a role for alternative secretion pathways for different proteins, as already proposed by Nogueira et al. (Nogueira et al., 2014). Following this logic, overexpressed βPS integrin would then also be trafficked through a different route from that of the endogenous βPS integrin, possibly because of higher expression levels or because of the presence of the Venus fused to the normal protein. Another reason for the sensitivity of Crb to loss of Tango1 may be that it is intensively recycled from the plasma membrane in tracheal cells and other tissues (Roeth et al., 2009; Schottenfeld-Roames et al., 2014; Sollier et al., 2015). We may therefore be seeing recycling Crb being trapped as a secondary consequence of the defects in the ER and Golgi system, whereas βPS integrin might remain in the plasma membrane once it has reached the cell surface, and therefore be able to gradually assembly there to near-normal levels even when it is partly blocked in its transit.

### Other secondary Tango1 cargos in fat body: interdependence of extracellular matrix proteins

Drosophila Tango1 was initially found to facilitate collagen secretion in the fat body. More recently, the accumulation of other non-bulky proteins at the ER in the absence of Tango1 has led to the proposal of two models to explain these results: One in which Tango1 regulates general secretion (Liu et al., 2017), and the second one where Tango1 is specialized on the secretion of ECM components (Liu et al., 2017; Tiwari et al., 2015), since loss of Tango1 leads to the accumulation of the ECM molecules SPARC and collagen (Tiwari et al., 2015). Our results suggest a third explanation, where cargo accumulation in the ER might not necessarily be a direct consequence of only the loss of Tango1. Instead, in addition to depending on Tango1, some proteins of the ECM appear also to depend on each other for their efficient secretion. This is the case for laminins LanB1 and LanB2, which require trimerization prior to exiting the ER, while LanA can be secreted as a monomer, (Hamill et al., 2009). Loss of collagen itself leads to the intracellular accumulation of ECM components in fat body cells, such as the laminins and SPARC. Conversely, SPARC is required for proper collagen and laminin secretion and assembly in the ECM (Martinek et al., 2008; Pastor-Pareja and Xu, 2011; Shahab et al., 2015). Furthermore, intricate biochemical interactions take place between ECM components (Kramer, 2005). Hence, due to the complex genetic and biochemical interactions between ECM components, the dependence of any one of them on Tango1 is difficult to determine without further biochemical evidence. The concept of interdependent protein transport from the ER as such is not new, as it has also been observed in other systems, for instance in immune complexes. During the assembly of T-cell receptor complexes and of IgM antibodies, subunits that are not assembled are retained in the ER and degraded (Call and Wucherpfennig, 2004; Geva and Schuldiner, 2014).

Nevertheless, our observations in glial cells, which express laminins but not collagen, allow us to at least partly separate these requirements. We find that laminins are accumulated due to general ER clogging and not because they rely on Tango1 for their export. This is based on our observations that once the protein causing the ER block is removed, laminin secretion can continue in the absence of Tango1. It is still unclear why glial cells can secrete laminins in the absence of collagen whereas fat body cells cannot, but presumably laminin secretion can be mediated by different, unidentified cargo receptors expressed in glial cells.

### A direct role for Tango1 in ER-Golgi organization

We found that Sec16 forms aberrant aggregates in cells lacking Tango1, as in mammalian cell lines (Saito et al., 2009), and that the number of Sec16 particles is reduced. Other studies have shown that Tango1 overexpression produces larger ERES (Liu et al., 2017), and that Tango1 and Sec16 depend on each other for localization to ERES (Maeda et al., 2017). In addition, as shown here and by others, lack of Tango1 also affects the distribution of Golgi markers (Bard et al., 2006; Liu et al., 2017; Santos et al., 2015). Thus Tango1 influences not only the trafficking of cargos, but also the morphology of the secretory system.

It had been suggested that the disorganization of ER and Golgi in cells lacking Tango1 might be an indirect consequence of the accumulation of Tango1 cargo (Saito et al., 2009). The work of Maeda et al. has provided a possible explanation for the molecular basis, and proposed that Tango1 makes general secretion more efficient, but it has not formally excluded the possibility that the primary cause for the observed defects is secretory protein overload. We have now shown that this is not the case: in the absence of Tango1 we still observe an aberrant ER and Golgi morphology even after we have removed the main primary substrates of Tango1 and thereby restored secretion of other molecules and prevented the ER stress response. The finding that Tango1-depleted cells have a functional secretory pathway in spite of the ER-Golgi disorganization was unexpected. Stress stimuli like amino acid starvation (but not ER stress response itself) lead to Sec16 translocation into Sec bodies and inhibition of protein secretion (Zacharogianni et al., 2014). However, uncoupling of ER-Golgi organization from functional secretion has also been observed in other contexts. Loss of Sec23 or Sec24-CD leads to KDEL appearing in aggregates of varying sizes and intensities similar to those we observe for Sec16 and for KDEL-RFP in cells lacking *tango1* (Norum et al., 2010). Also GM130 is reduced in Sec23 mutant embryos. However, these embryos do not show generalized secretion defects and also do not affect the functionality of the Golgi apparatus, as determined by glycosylation status of membrane proteins (Norum et al., 2010).

Thus, Tango1 appears to have an important structural function in coordinating the organization of the ER and the Golgi apparatus, and this in turn may enhance vesicle trafficking. This fits with the role of Tango1 in recruiting ERGIC membranes to the ERES (Santos et al., 2015), and also with the effects of loss of Tango1 in the distribution of ER and Golgi markers (as shown here and by others). Lavieu et al. have proposed that the ER and Golgi in insects, which unlike in mammalian cells is not centralized but spread throughout the cytoplasm, is less efficient for secretion of bulky cargo than mammalian cells that can accommodate and transport it more efficiently through the Golgi ribbon (Lavieu et al., 2014). This difference could explain why *tango1* knockout mice seem to have only collagen secretion defects and die only as neonates (Wilson et al., 2011). However, a complete blockage of the ER might also be prevented by the activity of other MIA3/cTAGE5 family homologs in mice. In mammalian cell culture experiments, even if loss of *tango1* affects secretion of HRP, the secretion of other overexpressed molecules like alkaline phosphatase is not affected. This could also be because of the presence of other MIA3/cTAGE5 family homologs. By contrast, because there are no other MIA3/cTAGE5 family proteins in Drosophila, loss of *tango1* may lead to the accumulation of a wider range of overexpressed proteins and more overt mutant phenotypes than in mammals.

## Acknowledgements

We thank S. Kraus for technical assistance, N. Jayanandanan for guidance in the early part of this work and S. De Renzis, P. Domingos, D. Gilmour, K. Roeper and members of the Leptin lab for helpful discussions. We are grateful to the Vienna Drosophila Resource Center, the Bloomington Drosophila Stock Center, the National Institute of Genetics Fly Stock Center, the Kyoto Drosophila Genetic Resource Center, the Drosophila Genomics Resource Center, and our colleagues M. Affolter, P. Domingos, S. Horne-Badovinac, C. Klämbt, E. Knust, S. Luschnig, C. Rabouille, C. Samakovlis, F. Schnorrer, G. Tanentzapf and B. Thompson for stocks and reagents. We thank the EMBL Advanced Light Microscopy Facility (ALMF) for continuous support and Zeiss for the support of the ALMF. We thank P. Bun for his support on image analysis pipelines. FlyBase was used throughout this work and is greatly appreciated.

The work was supported through funding from EMBO, EMBL, DFG grant LE 546/7-1 and the NRW Graduate School for Genetics and Functional Genomics. LDRB was funded by the EMBL Interdisciplinary Postdoctoral Programme under Marie Curie Actions.

## Author contributions

MB identified and mapped the 2L3443 mutation. SS generated the UAS-Tango1-GFP, and UAS-βPS-Integrin-Venus lines. Experiments were designed by LDRB, SS and ML, and performed by LDRB and SS. LDRB and ML wrote the paper. All authors have read and edited the manuscript.

